# Tracking butterfly flight in the field from an unmanned aerial vehicle (UAV): a methodological proof of principle

**DOI:** 10.1101/2024.07.17.603869

**Authors:** Emmanuel de Margerie, Kyra Monmasson

## Abstract

Tracking and understanding the movements of animals in the wild is a fast-growing area of research, known as *movement ecology*. However, tracking small animals such as flying insects, which cannot easily carry an electronic tag, remains challenging as existing field methods are costly either in terms of equipment or tracking effort (e.g. VHF radio-tracking, scanning harmonic radar). Here we attempted to record the movements of free-flying butterflies from an unmanned aerial vehicle (UAV), maintaining a static position in the sky and recording video vertically downwards. With an appropriate flight height and image filtering algorithm, we recorded 166 flight tracks of *Pieris* butterflies (*P. brassicae* and *P. rapae*), with a median tracking length of 40 m (median flight duration 13 s), and a high temporal resolution of 30 positions per second. Average flight direction varied significantly over the course of the flying season, from a northward azimuth in June and early July, to a southward azimuth in September, congruent with a trans-generational migratory behaviour that has previously been documented by field observations or experiments in flight cages. In addition, UAV imagery unlocks the possibility to measure high-resolution flight movement patterns (e.g. path tortuosity and transverse oscillations), which will possibly help understand perceptual and locomotor mechanisms underlying spatial behaviour. We explore the technical details associated with UAV tracking methodology, and discuss its limitations, in particular the biases associated with a 2D projection of 3D flight movements, the limited spatial scale, and the difficulty to distinguish between visually similar species, such as *P. brassicae* and *P. rapae*.

## Introduction

Tracking the movements of an animal in the wild provides insights on its ecology, such as habitat use (e.g. Da Silveira et al 2016), dispersal and migration behaviours (Whitfield et al. 2024, Rotics et al. 2016). When the movement data are detailed enough, both spatially and temporally, it is also possible to extract biological information on the animal’s locomotor biomechanics (Sherub et al. 2016, Hedrick et al. 2018, Ruaux et al. 2023), its spatial search strategies (Shepard et al. 2011, Hernandez-Pilego et al. 2017, de Margerie et al. 2018), and more broadly the perceptual and cognitive processes involved in movement (Kashetsky et al. 2021). All of these biological and ecological aspects of animal movement have been advantageously integrated into a synthetic movement ecology framework (Nathan et al. 2008, Abrahms et al. 2017, Joo et al. 2022), that is progressively articulated with other major fields of biological research (e.g. community ecology: Schlägel et al. 2020; animal physiology: Hetem et al. 2025).

However, not all animal species are easy to track. Large-enough animals (approx. > 100 g) can carry GPS receivers (Cagnacci et al 2010, Wilmers et al 2015), which are now often enhanced with additional sensors (accelerometers, barometers, cameras) to better infer the animal’s behaviour along its route (Kays et al 2015, Joo et al. 2022). To track the movements of smaller species such as flying insects, tracking tags need to be much lighter, and alternative technologies to GPS are employed. Beacons emitting simple VHF radio beeps have been used successfully for decades to track the movements of flying and non-flying insects (Kissling et al. 2014). Yet, VHF radio-tracking involves following the animal with one or multiple antennae (or deploying a fixed antennae array; e.g. Knight et al. 2019), and the spatio-temporal resolution is inferior to GPS tracking. Even lighter, passive tags (of a few mg, without any battery) can be used to track the movements of flying insects (Ovaskainen et al. 2008, Lihoreau et al. 2012, Maggiora et al. 2019). The downside of these passive tags is that a scanning harmonic radar (SHR) - a heavy and expensive device - has to be deployed in the field. The tracking range of SHR is near 1 km, and the temporal resolution of the data is one position every 3 s (Ovaskainen et al. 2008). Other shorter-range, passive-tag tracking systems also exist (portable harmonic radar, RFID tags; see Kissling et al. 2014, Batsleer et al. 2020, Rhodes et al. 2022 for reviews).

For all animal-borne tracking systems, whether GPS receivers or other types of tags, the impact of the tag on the animal’s movements is a matter of concern. The carried mass, but also the drag, the position, or the method of attachment of the tag must be carefully considered. Impact studies are often necessary, both from an ethics point of view and in terms of the reliability of the collected movement data (Wilson & McMahon 2006, Batsleer et al. 2020).

There are other routes to track flying insects, without having to place tags on animals. First, the classic mark-recapture method allows to demonstrate the movement of an individual from one point to another, and was thus used for studying insect dispersal and migration (e.g. butterflies; Chowdhury et al. 2021, 2022). Trapping techniques along flight routes, sometimes combined with isotope analyses to determine the geographical origin of individuals, can also provide information on large scale movements (e.g. dragonflies; Knoblauch et al. 2021, Oelman et al. 2023). At a finer, local scale, if the insect flies slowly enough, in an open environment where it remains visible, it can be followed by foot and its passed positions can be drawn on a map (Brussard & Ehrlich 1970) or recorded by a GPS receiver carried by the observer (Delattre et al. 2013, Fernandez et al. 2016). Tracking butterflies crossing a body of water from a small boat is another possible technique (Srygley & Oliveira 2001). Also, when flying insects move in swarms of many individuals, radars (weather surveillance radars or smaller scale biological radars) can be used to detect and measure the direction of these flights (e.g. Stefanescu et al. 2013). With biological radars, individual movement variables such as flight direction, height and ground speed can be extracted (Bauer et al. 2024).

To complement these “tagless” tracking approaches, sometimes called “passive sensing” (Rhodes et al. 2022), we wondered whether flying insects could be tracked in video images filmed from a UAV (Unmanned Air Vehicle, or drone) positioned above them in the sky.

Image-based tracking from ground-based cameras is a known technique to reconstruct the 2D and 3D trajectories of animals, and in particular flying insects, with a high sampling frequency (> 1 Hz). It can be used in laboratory settings (e.g. Lihoreau et al. 2016a), semi-natural outdoor insectaries (Kitamura & Imafuku 2015, Le Roy et al. 2021, Kleckova et al. 2024) or even natural environments (Stürzl et al. 2016, Jackson et al. 2016). Most often used to address biological questions related to perception, cognition or locomotion, image-based tracking techniques, which are less invasive, are also attracting growing interest in ecology (Dell et al. 2014). Hybrid techniques tracking tagged animals in videos are also being developed (Crall et al. 2015, Walter et al. 2021).

On the other hand, in the last decade, commercial UAVs have greatly improved in terms of compactness, stability and image resolution, while decreasing in cost, making them valuable tools for wildlife inventory and conservation (Wang et al. 2019, Charbonneau & Lemaître 2021). Tracking animal movements using videos recorded from a UAV is a next logical step, and it has already been achieved for a variety of large species (e.g. reef shark: Rieucau et al. 2018, wild dog: Haalck et al. 2020, zebras and geladas: Koger et al. 2023). For flying insects, UAV-image-based tracking has been proposed previously (Ivosevic et al. 2017), but remains to be tested and validated. Most recently, Vo-Doan et al. (2024) successfully tracked a honey bee from a special UAV-borne optical system (Fast Lock-On), but this technique requires that the insect carries a reflective marker.

Here we explore the validity of a UAV as a platform for remote, passive observation of the movements of untagged insects in flight. The motivation for this exploration is the many potential benefits of such an approach, namely (1) non-invasiveness, i.e. tracking of animals moving freely, without tags and associated capture procedures, (2) in an open natural environment, over distances greater than in an insectary, (3) with a spatio-temporal resolution superior to VHF, SHR or GPS tracking. If this type of fine movement data in natural conditions can be collected easily, it could be very useful for bridging the gap between laboratory studies on perceptual, cognitive and locomotor mechanisms, and movement patterns observed in the natural environment at the local scale (daily routine movements) or beyond (insect dispersal or migration).

To begin this methodological exploration with a relatively simple case, we have focused on *Pieris* butterfly species (large white *Pieris brassicae* and small white *Pieris rapae*) because of their relatively slow flight, good visibility and abundance in the field. We limited the present study to the simple situation of a single, static UAV in the sky, filming vertically downwards (Fig. 1A) to reconstruct flight trajectories in only 2 horizontal dimensions. We also chose to record flights accross areas with low ecological resources, which are likely to promote simple directed movements rather than highly tortuous, resource-searching movements (Schtickzelle et al. 2007, Fernandez et al. 2016, Schlagel et al. 2020).

**Figure 1.**
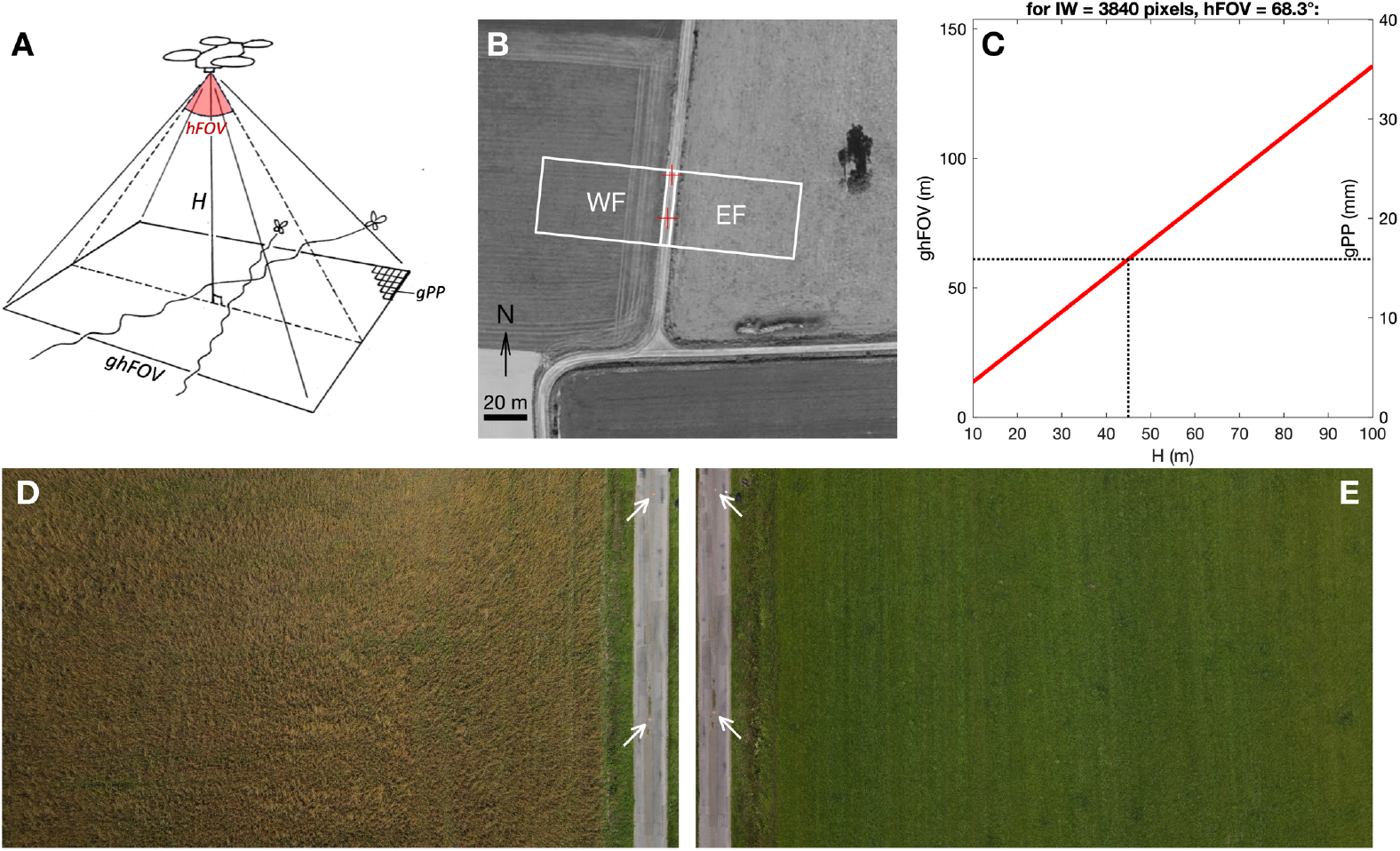
UAV field of view (FOV). **(A)** Video recording geometry, showing the influence of UAV flight height (*H*) and camera angular horizontal FOV (*hFOV*) on ground horizontal FOV (*ghFOV*) and ground pixel pitch (*gPP*). **(B)** Aerial view of the study site, showing FOV over western (WF) and eastern field (EF). *Source for aerial photography: geoportail*.*gouv*.*fr* **(C)** Variation of *ghFOV* and *gPP*, as functions of *H*. These linear relationships depend on camera angular *hFOV* and recorded image width *IW* (see equations 1-2). **(D-E)** Examples of video frames recorded on July 8^th^ 2021, over WF and EF, respectively. White arrows identify 2 reference crosses painted 20 m apart on the road, and used for image scaling.

## Material and Methods

This section describes the general methods we used to film butterflies in the field, reconstruct their flight trajectories and describe their movements. We later explored and validated these data using diverse specific methods and statistics, which are detailed at the start of each results sub-section, for ease of reading.

### Study site

We recorded butterfly flights from June to September 2021, in an agricultural area near the city of Rennes, France (coordinates 48.105444, −1.560029). The local landscape is covered with cultivated fields, few tree patches, small roads and farms, with the closest urbanized area situated 1.7 km away. We filmed butterflies flying above 2 fields situated on each side of a narrow road (Fig.1B). The western field (WF) contained an organic mixed crop (mainly wheat, fava and lucerne) which was harvested in August, whereas the eastern field (EF) contained forage grass, which was cut in June and August. We chose to record butterflies above two different fields (filmed in alternance) to test our image filtering method above various backgrounds, and also control whether the flight trajectories could be influenced by ground vegetation: while EF crop was devoid of any identified ecological ressource, WF contained a few nectar-bearing flowers that might attract butterflies, and hence influence their flight trajectory.

On the central road were the UAV takeoff/landing area, the UAV pilot (EDM), the technical assistant (KM), and an ultrasonic anemometer (Gill Maximet 501) recording wind speed and the direction from which the wind originates every second, at 2 m height. We painted two permanent red crosses 20 m apart along the central road (Fig.1D, E), as a reference line segment for positioning the UAV, and later for scaling the video frames. The ground slope in the recorded area was less than 2°.

### UAV video recording

We used a Mavic Air 2 UAV (DJI, Nanshan, Shenzhen, China), which is a small commercial quadricopter (takeoff weight 570 g, retail price ∼1000 € in 2021). This UAV has a CMOS sensor (6.4 × 4.8 mm) which can record 3840 × 2160 pixel videos (i.e. “4K” images, with 16:9 aspect ratio). According to the UAV manual, the camera lens has an f/2.8 aperture and an 84° field of view. As most recent UAVs can record with various aspect ratios and resolution levels (which may involve sensor cropping, i.e. digital zoom), we prefered to measure FOV in the lab, by placing the UAV camera at a known distance from a wall, and measuring the horizontal distance along the wall that is effectively included in the UAV camera image. The “horizontal” FOV (*hFOV*, i.e. along image width), was measured at 68.3° in the default “4K wide” recording mode, that we used throughout the present study. This angular *hFOV* value was used to choose a flight height. We computed the horizontal field of view on the ground (*ghFOV*, in meters) when the UAV camera aims vertically downwards:

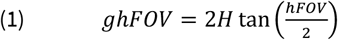

where *H* is the UAV height (m), and *hFOV* is the horizontal angular FOV (°).

Proportional to *ghFOV* is the corresponding pixel pitch on the ground, i.e. the distance on the ground covered by a single pixel side (*gPP*, m):

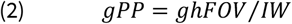

where *IW* is the video image width, in pixels.

The total recorded ground area (*gaFOV*, m^2^), can also be of interest:

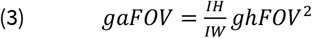

where *IH* is the video image height, in pixels.

Based on these relations, we chose a 45 m UAV flight height, which covers a *ghFOV* of 61 m (*gaFOV* = 2096 m^2^), and corresponds to a *gPP* of 16 mm (Fig. 1C). This pixel pitch value was voluntarily chosen at a fraction of the body size of the *Pieris* species we wanted to track, which have a forewing length around 30 mm.

We performed preliminary tests in the field that confirmed that at *H* = 45 m, *Pieris* butterflies flying near the ground were projected in recorded images as pixel “blobs” with an area around 10 pixels, which is large enough to be reliably tracked from frame to frame (see Fig. 2A, C). Higher camera height would allow larger FOV area on the ground, and hence longer tracking durations, but automatically tracking smaller blobs would become less reliable.

**Figure 2.**
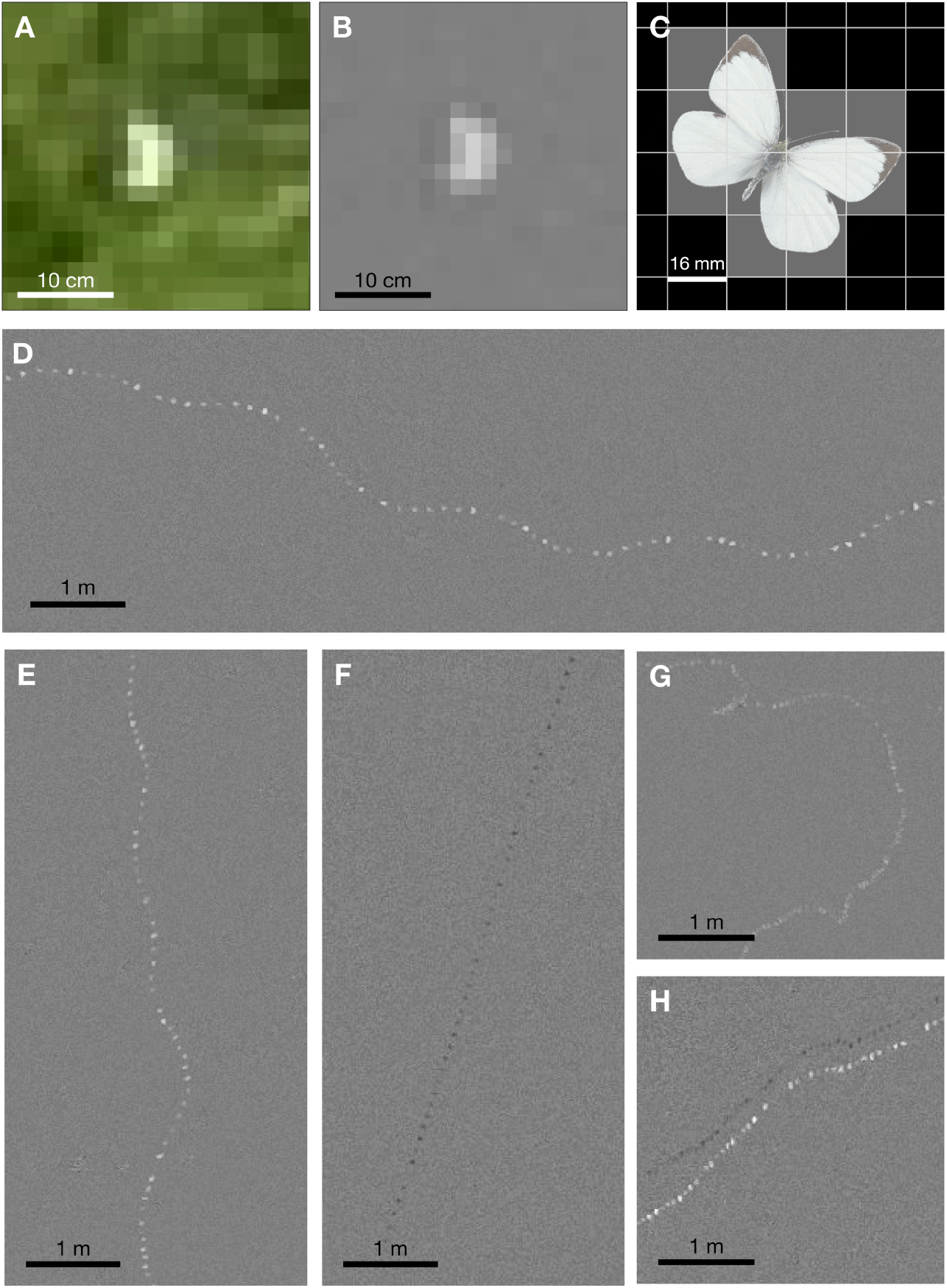
Video frame filtering. **(A)** Magnified view of an original video frame, containing a *Pieris brassicae* image (as identified after butterfly capture). **(B)** Same view after applying the blink filter (i.e. *b* frame). **(C)** A hypothetical 30 mm forewing length *Pieris*, with wings fully stretched, projected onto a pixel grid with a 16 mm *gPP*. If all partially covered pixels appear brighter, the pixel blob area would be around 12 pixels. **(D)** Merging successive *b* frames reveals *P. brassicae* flight trace in a synthetic, diachronous image (i.e. *B* image). **(E)** Example trace of *Pieris rapae*. **(F)** Example trace of a dark-coloured species, either *Vanessa atalanta* or *Aglais io* (contrast ×2.0). **(G)** Example trace of *Melanargia galathea*. **(H)** Example trace where the shadow of a *Pieris* butterfly is also visible (contrast ×1.5).

Note that at *H* = 45m, the butterflies are not visible to the UAV pilot through the live video feedback on the UAV controller screen. The video feedback has lower resolution (*IH* = 720 or 1080 pixels) than the recorded video, and the controler screen (Ipad Mini 5, Apple, Cupertino, USA) also has limited magnification and contrast in outdoor conditions. Butterflies in UAV footage were only detectable *a posteriori*, when playing recorded videos at full resolution on a computer screen in the lab.

All videos were recorded at “30 fps” (29.97 video frames per second), which in 4K resolution produced video data at a rate of 13 MB/s. The UAV firmware automatically cut videos lasting more than 5 min in multiple 4 GB files, which can be stitched in post-processing, but with the loss of one video frame between files (this may depend on the UAV model). For simplicity, we decided to keep each video duration below 5 min. Recording at 60 fps was another possible option, but with a 15 MB/s data rate, this implied less data collected per video frame. We sticked to 30 fps, aiming to obtain the best possible image quality per video frame.

Constant, manual exposure was tested initially, but proved unpractical as the ground luminance could vary by several exposure values (Ev) when cloud shadows crossed the FOV. Hence, we used auto-exposure with an exposure compensation of −0.7 to −1.3 Ev, as it improved the contrast between the white butterflies and the background.

### Time distribution of UAV flights

In order to distribute observations accross the *Pieris* flight season (which in France can span from April to October; Lafranchis et al. 2015), we chose to collect videos once a week, over the months of June, July and September 2021. Each week, we chose a day that was favorable for butterfly flight, i.e. with a weather forecast as warm, sunny and not very windy as possible. We went on site in the afternoon (14:00-16:30). After takeoff, the UAV was positioned at *H* = 45 m above the EF. The UAV camera was tilted to a vertically-downward position, and the pilot used the video feedback to place the scale points on the road to the left margin of the FOV (see Fig. 1E). The pilot started the video recording, and the UAV then relied on its own sensors (GPS, altimeter, etc.) to maintain its position without any pilot input, for about 4 min 30 s. Then the video recording was stopped, the UAV was relocated to the WF, height and scale alignment (now on the right margin of the FOV, see Fig. 1D) were checked, and another 4 min 30 s video was recorded. Repeating these steps, we could record a second EF video and a second WF video before the UAV battery dropped to 20-30% of capacity, inciting landing the UAV and swapping its battery. With this sequence, each UAV battery (rated at 11.6 V, 3500 mAh, 40.4 Wh) allowed to record about 18 min of video. As we used 3 fully charged batteries per field session, we were able to collect about 1 hour of video per field session. Weather allowing, 3 field sessions could take place in June (1^st^, 9^th^, 15^th^), 4 field sessions in July (1^st^, 8^th^, 15^th^, 22^nd^) and 4 in September (6^th^, 16^th^, 22^nd^, 29^th^), for a total video duration of 10.7 hours (48.8% over WF, 51.2% over EF).

### Video processing

In order to automate the tracking of butterflies in videos, pixels corresponding to the butterfly should be easily distinguished from the background. In the present case, *Pieris* butterflies appear in the raw video as blobs of bright pixels, but the background formed by the vegetation also has multiple bright areas (Fig. 2A), which rules out a simple detection of the butterfly by thresholding the raw video luminance levels. In addition, as the UAV is not perfectly static, and the wind can cause vegetation on the ground to move slowly, the background is moving, which does not favour background subtraction approaches (Piccardi 2004). We found a solution to this issue by designing a custom filter that selects the pixels that blink in the video: when a butterfly passes over an area, the pixels in that area become brighter for only 1 frame, and then revert to the background luminance. To apply this filter, we first transformed the RGB video frames into greyscale. Each pixel then has a single luminance value (*v*) in the range [0, 255]. Then the “blink” filter script performs the following calculations:

Pixel value variations from current frame (t) to next (t+1) and previous (t-1) video frames:

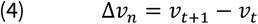

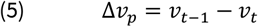

If both variations have the same sign, there was a luminance peak (positive or negative), and a blink value (*b*) is computed. On the contrary, if variations have different signs, there was no blinking, only monotonous pixel value variation.

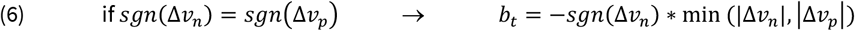

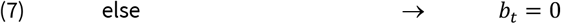

Note that *b* can be positive (bright blob passing over darker background) or negative (dark blob passing over brighter background).

Interestingly, when going through a series of successive video frames, keeping memory of extreme *b* values for each pixel (notated *B*) can be used to reveal flight trajectories as a series of blobs, in a synthetic, diachronous image.

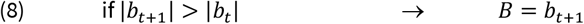

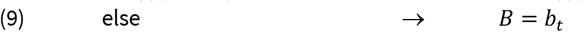

For easy display of *b* frames (and *B* synthetic images), pixel blink values are rescaled from [-255, 255] to [0, 255].

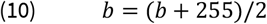

As a result, background appears as medium grey (*b* = 128), with blinking pixels as lighter or darker gray blobs. Fig. 2 shows example results of the blink filter.

We used the blink filter in two processing steps. For the initial exploration of our videos, we cut each video into 20 s bouts, and generated a single *B* image summarising each bout. This enabled us to quickly detect which video bouts contained butterfly tracks. On this basis, the FOV entry and exit times of each butterfly were precisely noted by playing the original videos on a large screen. Based on these time limits, we then generated for each butterfly track a new greyscale, uncompressed video file, containing the series of *b* frames. This filtered greyscale video was then used to automatically track the blobs.

We used DLTdv8 (Hedrick 2008) to extract blob coordinates (in pixels) in successive video frames. The blob was manually digitized (i.e. mouse clicked) in the first few frames, and then DLTdv8 uses a Kalman filter and 2D cross-correlation to find the blob in following frames (without thresholding). As blob size and contrast can vary, and filtered *b* frames still contain some noise in the background, the automatic tracking process needed human supervision and frequent manual corrections, but with a convenient user interface to navigate through video frames, DLTdv8 offered vast time savings compared to a fully manual digitization. The average time spent on screen to process the videos was approximately 15 min per track. There are numerous alternative options to DLTdv8 for tracking blobs through video frames (e.g. Sridhar et al. 2019, Lauer et al. 2022, Chiara & Kim 2023). Regardless of the tracking software used, starting from the blink-filtered video will help solve the natural, moving background issue.

For scaling the butterfly track to real-world coordinates, we measured pixel coordinates of the two painted reference crosses in one video frame (at mid-duration of the track), and fitted a geometrical transformation (combining rotation, scaling and translation; *fitgeotrans* function in Matlab) that resulted in (0, 0) and (0, −20) coordinates in real-world meters. This same transformation was then applied to the whole butterfly coordinate series, transforming coordinates in pixels to meters. We had measured with a compass in the field that the central road had a 6°E azimuth. We checked and refined this value in a GIS software (https://www.geoportail.gouv.fr), and thus applied a 5.83° CW rotation to all butterfly coordinates, so that in all graphical representations, y axis has a 0°, northward azimuth.

### Track selection

When we explored the *B* images synthesizing our videos, we mainly found flight traces of *Pieris* butterflies (large white *P. brassicae*, small white *P. rapae*), appearing as clearly visible white dotted traces (fig. 2D, E). These two species were also the most easily observed from the ground during the field sessions. We also found a few flight traces of other species, that we had observed in the field, such as the marbled white (*Melanargia galathea*), the red admiral (*Vanessa atalanta*) or the peacock (*Aglais io*). However, as these traces were less frequent, and usually barely visible and discontinuous (Fig. 2F, G), we did not analyse them further.

For some *Pieris* traces, the shadow of the butterfly formed a second, dark trace (Fig. 2H), which may potentially be used to determine the height of the butterfly relative to the ground (knowing the associated sun elevation angle). These shadows were only visible on almost bare ground (September videos, after crop harvest), and thus concerned a small number of *Pieris* traces. We have therefore not used this additional source of 3D data in the present study.

Among the *Pieris* tracks, we applied the following exclusion criteria:

- A 10 m wide strip, containing the road and the adjacent flowered ditches, was excluded from the analysis, so that each field (WF and EF) included a homogeneous area in terms of ground vegetation.
- Many very brief tracks (< 5 s), often crossing just one corner of the FOV, were considered less informative and excluded from analysis.
- We chose to focus on butterflies in continuous movement: Tracks marking one or more stops while crossing the FOV were excluded (a stop being defined as remaining for at least 1 s within a radius of 0.1 m). These tracks with stops were in the minority (∼ 1 out of 6 tracks).
- Tracks in which the butterfly interacted with another individual (e.g. flight inflection towards another butterfly, chases) were also excluded (∼ 1 out of 13 tracks).

In the end, our sample comprised *N* = 166 *Pieris* flight tracks. Despite our constant sampling efforts, these were not evenly distributed through the season: we recorded 12 tracks in June, 125 in July and 29 in September. Taking all these tracks together, 70524 butterfly positions were recorded, representing a total of 39 min of flight time, and a flight distance of 7.4 km.

When the tracks contained missing positions (caused by the absence of blob in some *b* frames), the (x, y) coordinates were linearly interpolated. These interpolated positions represented 2.2 % of the dataset (1544 positions). The interpolated positions were rarely contiguous, and the longest interpolated segment represented 7 successive positions (i.e. 0.23 s).

### Track descriptive variables

For this pilot study, we computed a small set of basic descriptive variables for each 2D track.

- **Track duration** is the time the butterfly remained within the FOV (road zone excluded).
- The change in 2D position from a frame to the next frame is named a step vector. **Track length** is computed as the sum of step vector norms.
- Step speed is equal to step vector norm divided by the elapsed time (i.e. 1/29.97 s). **Average speed** is the arithmetic mean of the series of step speed values. It indicates how fast, on average, the butterfly flew along its 2D track. Average speed is also equal to track length divided by track duration. Note that average speed is a ground speed, not an air speed.
- A “beeline” vector is defined as the vector from the first to the last recorded position of the butterfly. The beeline vector is equal to the vectorial sum of step vectors, and hence represents the butterfly’s resultant, directed movement across the FOV. **Beeline azimuth** was the direction of the beeline vector, a circular variable in the interval [0°, 360°[, 0° corresponding to a northward azimuth.
- Track **straightness** is computed as the ratio of the beeline vector norm to the track length. Straightness value is in the interval [0, 1]: 0 indicates that the butterfly performed a loop (i.e. had the same entry and exit positions), while 1 indicates a perfectly straight flight path. Straightness is inversely related to path tortuosity. Track straightness is also known as “Net to Gross Displacement Ratio” (NGDR, Buskey 1984).
- **Beeline speed** is the norm of the beeline vector divided by track duration. It reflects how fast the butterfly, on average, progressed in its directed movement. Beeline speed is also equal to average speed multiplied by straightness. Note that beeline speed is also a ground speed, not an air speed.
- **Wind speed** and **direction** for each track were computed from the vectorial sum of the n wind vectors recorded during track duration, divided by n.

### Processing and statistics software

Pixel blob tracking in videos was performed using DLTdv8 (Hedrick 2008; https://biomech.web.unc.edu/dltdv/). Other analyses, from video processing to statistics, were performed using Matlab R2018b (The MathWorks, Natick, MA, USA). We used the CircStat2012a toolbox for circular statistics (Berens 2009). For a small number of captured butterflies, forewing length measurement from field photographs were performed with ImageJ V1.54g (http://imagej.org).

## Results

### Track general description

Fig. 3 shows the reconstructed 2D tracks, and associated variable distributions. We collected 66 *Pieris* flight tracks over WF, and 100 tracks over EF. Tracks had a median duration of 12.8 s (range 5.1 to 56.5 s, Fig. 3B), for a median track length of 40.0 m (13.4 to 134.9 m, Fig. 3C). The median value for average speed was 3.3 m.s^-1^ (1.9 to 9.2 m.s^-1^, Fig. 3D), and 2.9 m.s^-1^ for beeline speed (0.2 to 9.1 m.s^-1^, Fig. 3F). Straightness distribution was strongly skewed towards straight tracks, with a median value of 0.93 (range 0.10 to 0.99, Fig. 3E). High straightness values imply similar values for average speed and beeline speed in most tracks.

**Figure 3.**
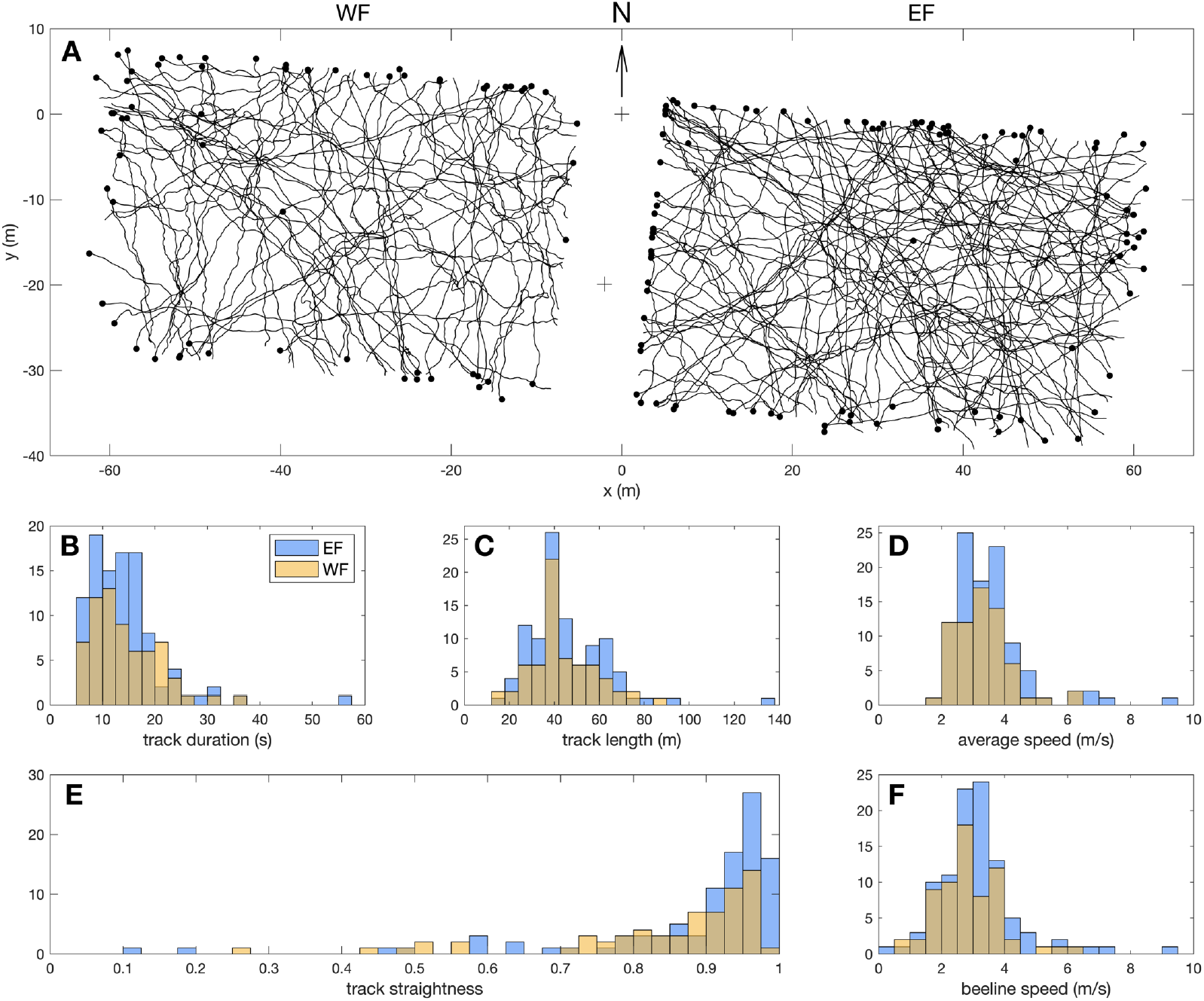
*Pieris* flight tracks description. **(A)** Reconstructed 2D tracks over WF and EF. Black dots indicate last position of each track. **(B-F)** Distributions of descriptive variables: (B) track duration, (C) track length, (D) average speed, (E) track straightness and (F) beeline speed.

We used two-sample Kolmogorov-Smirnov (KS) tests to assess distribution differences between WF and EF tracks, and found no significant difference for track duration (*D*_*(66,100)*_ = 0.10, *p* = 0.78), track length (*D*_*(66,100)*_ = 0.09, *p* = 0.86), average speed (*D*_*(66,100)*_ = 0.09, *p* = 0.86), or beeline speed (*D*_*(66,100)*_ = 0.18, *p* = 0.15). However, there was a significant difference between straightness distribution over WF and EF (*D*_*(66,100)*_ = 0.26, *p* = 0.009), with straightness for EF tracks being even more skewed towards 1 (median 0.94) than WF tracks (median 0.90; see Fig. 3E).

Beyond average speed and straightness, the 30 fps butterfly flight tracks extracted from UAV videos also contained fine-scale instantaneous information about flight speed and azimuth variation along the tracks. For example, Fig. 4 shows a track segment that depicts interesting movement patterns, in the form of meter-scale, sub-second transverse oscillations along the flight path.

**Figure 4.**
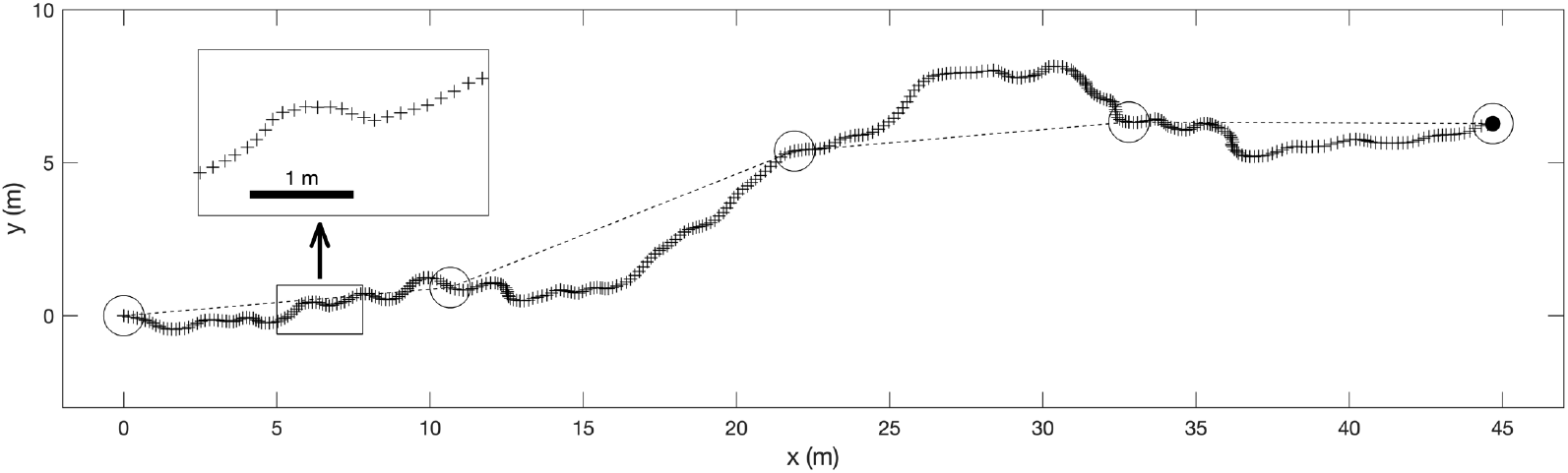
Benefit of 30 Hz positional data. A 12 s flight track segment, reconstructed with the present method, showing all recorded 2D positions (+, 361 in total). The insert shows a magnified view. The 5 circles represent the positions that could have been recorded with a SHR tracking system, that has a 90-fold lower sampling frequency (1 position every 3s.). Note that on the other hand, SHR benefits from a larger FOV, and hence longer tracking durations (see discussion).

### Controlling UAV stability

During video recording, the UAV was not perfectly static, and whether this could have an effect on the reconstructed flight tracks was an important issue. For a small random subset of tracking videos (*N* = 10), we digitized the two reference points painted on the road, describing a 20 m reference segment (RS), not only at mid-duration, but on every video frame (using automatic tracking in DLTdv8), which allowed to monitor how the RS was transformed throughout the duration of tracking, due to UAV movements. We assumed that a combination of rotation, scaling and translation could affect the RS projection. Using the *fitgeotrans* function in Matlab, we obtained the geometric transformation matrix from the first frame’s RS to each following frame, which allowed to monitor rotation, scaling and translation movement components separately.

Fig. 5 shows that the RS projected image was indeed affected by a combination of geometric transformations through time. Over the investigated tracking durations (13 to 40 s), rotation of the RS could attain 0.26° (Fig. 5A), and was on average 0.07° (root mean square, RMS). Scaling variation could attain ± 1.13 % (0.36 % RMS; Fig. 5C). Horizontal (x) or vertical (y) translation of the RS image (Fig. 5B) could reach ± 11.2 pixels (3.6 pixels RMS), which represents less than 0.3 % of image width, or 0.18 m when projected on the ground. The amounts of transformation usually did not grow monotonically through time, reflecting that in the absence of pilot input, the UAV does not simply drift away from its initial position, but uses inputs from its onboard sensors to try and maintain position and azimuth. Digitizing the RS only once at track mid-duration - i.e. assuming that the UAV is fully static during each track - resulted in a maximal 0.28 m error (0.08 m RMS) on the butterfly reconstructed 2D position (Fig. 5D). For the present work, we considered these levels of error to be acceptable, which is why we used the single RS digitizing method for all the remaining tracks.

**Figure 5.**
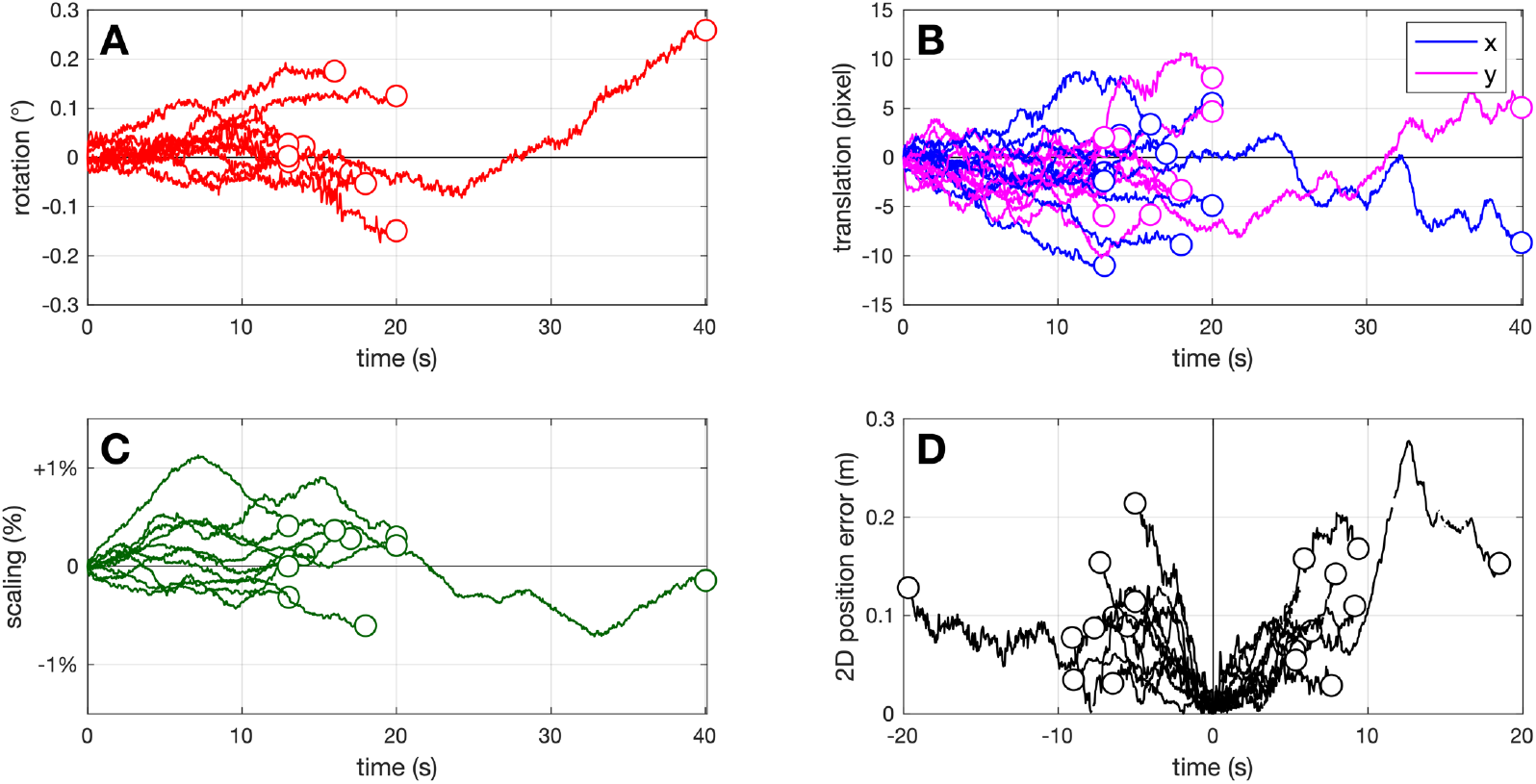
UAV movements. Reference segment transformation through time for 10 tracking videos, decomposed into **(A)** rotation, **(B)** translation and **(C)** scaling components. **(D)** Resulting 2D error on the butterfly 2D position, when only a single digitization of the RS is performed at mid-duration.

### Controlling butterfly flight height

With the present recording geometry, the flight tracks of all butterflies passing through the pyramidal FOV (Fig. 1A), regardless of their flight height, are projected on a single sensor plane. We reconstructed these tracks in 2D by assuming that the butterflies moved in the same plane as our reference 20 m segment, i.e. flew at ground level. How this simplification departed from the 3D reality needed investigation. We were especially interested in the possibility that some butterflies might have crossed the FOV at a significant height (10-40 m), which would result in vastly overestimated flight speeds in our ground-projected reconstructions.

During field work, we visually monitored the fields under the UAV, and noted each time we saw a *Pieris* butterfly passing at a low height (defined as below the observer visual horizon, i.e. less than ∼2 m above the ground). By comparing our field notes with the timestamps of the reconstructed tracks, *N* = 45 out of 166 tracks (i.e. 27 %) corresponded to visually-detected butterflies flying at low height (LH group). The remaining tracks were qualified as “unknown height” (UH group, *N* = 121), and may correspond either to high-flying butterflies (visually undetected because of low contrast against the sky), or to low-flying butterflies that remained undetected (because the observer’s attention was regularly directed at the UAV rather than at the ground).

We compared the beeline speeds of the low-height (LH) and unknown-height (UH) groups. Both groups had very similar speed distributions (Fig. 6A), as confirmed by statistical tests : LH (2.9 ± 1.0 m.s^-1^, mean ± SD) and UH tracks (3.1 ± 1.3 m.s^-1^) did not significantly differ for mean speed in a *t*-test (*t*_(164)_ = 0.84, *p* = 0.40), and were not drawn from different distributions according to a KS test (*D*_(45,121)_ = 0.11, *p* = 0.79).

**Figure 6.**
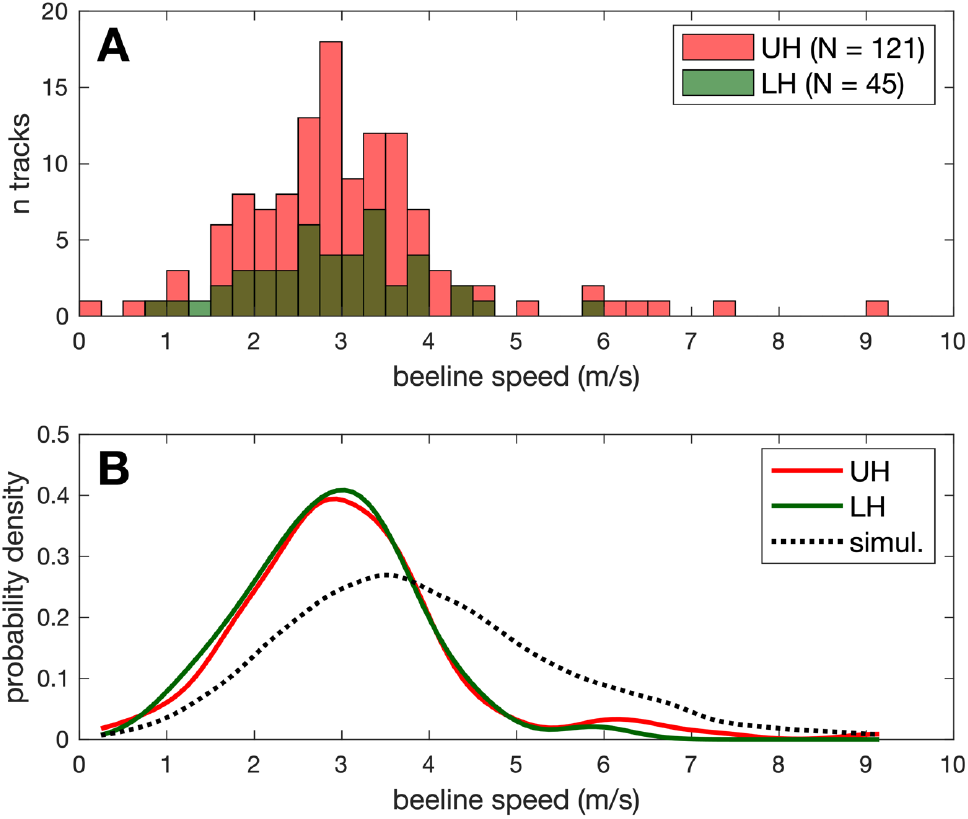
Speed distribution and flight height. **(A)** Ground beeline speed distribution observed for low height (LH) and unknown height (UH) butterfly tracks. **(B)** Kernel probability density estimation for LH and UH tracks, and for simulated tracks with uniform height distribution (see text).

In order to visualize what speed distribution would be obtained for butterflies flying far from the ground, we simulated 10^5^ straight horizontal tracks, with flight speeds sampled from a normal distribution copied form the LH group (2.9 ± 1.0 m.s^-1^), but crossing the FOV at heights uniformly distributed between 0 and 45 m. Simulated tracks crossing the FOV, and with a track duration of at least 5 s (to simulate comparable track selection), had a ground-projected speed distribution that was flatter and shifted towards high (i.e. overestimated) speeds when compared to both LH and UH tracks (see Fig. 6B). This further suggests that the UH group in our track sample mainly comprises low-flying butterflies that went visually undetected in the field. Still, we cannot exclude that a small number of tracks in the UH group (on the right tail of speed distribution, e.g. with reconstructed speed > 6 m/s) might correspond to high-flying butterflies. Using the pixel blob area as a proxy to measure flight height was not considered a valid option, as the blob area can vary considerably even for a single individual flying at low height (see next section).

### Exploring blob area as a specific signature

We explored the possibility to discriminate *P. brassicae* from *P. rapae* tracks, based on pixel blob area in the videos. During field work we captured a small number (*N* = 8) of *Pieris* individuals that had just passed through the UAV’s FOV. Each individual, captured with a butterfly net, was briefly placed in a thin transparent box with a grid-patterned back, photographed with its wings stretched for later identification and size measurement, and then immediately released. Back in the lab, we measured each individual’s forewing length (from wing base to wing tip, Van Hook et al. 2012) using ImageJ. The UAV video recordings corresponding to these individuals were filtered and digitized with the same methods as previously described, but were later re-analysed to measure the blob size in each filtered video frame. As a simple approach, the blob was defined as connected pixels with grey level *b* > 138, i.e. departing by more than 10 grey level from the mean background grey of 128. Sometimes the simple thresholding approach detected no blob, but this method still allowed to obtain many (269 to 654) blob area values per individual, that could be compared to the animal’s real size. Note that 4 out of the 8 tracks used for the present blob area analysis were not part of the final flight track sample (*N* = 166), because these captured individuals had flown near the road, or performed stops along their flight (see Track selection section).

Five captured individuals were identified as *P. rapae* (2 females + 3 males), with forewing length ranging from 22.8 to 28.0 mm. The 3 other captured individuals were *P. brassicae* (2 females + 1 male), with longer forewing, ranging from 31.2 to 36.7 mm. These values were in line with the literature, with the largest *P. rapae* individuals being close in size to the smallest *P. brassicae* specimens (Cook et al. 2022).

Fig. 7 shows the blob area distributions observed for all 8 captured individuals. When considering only the median blob area for each individual, it was positively correlated to the forewing length of the animal (Spearman rank correlation, *r*_(6)_ = 0.73, *p* = 0.047). However, the relationship was not monotonically increasing (*r* < 1), and there was extensive overlap between blob area distributions. In other words, a smaller butterfly could often project as a larger pixel blob than a larger butterfly, depending on the compared video frames. As a result, we considered unreliable to use recorded pixel blob areas as a direct mean to discriminate *P. rapae* and *P. brassicae* flight tracks in the present work.

**Figure 7.**
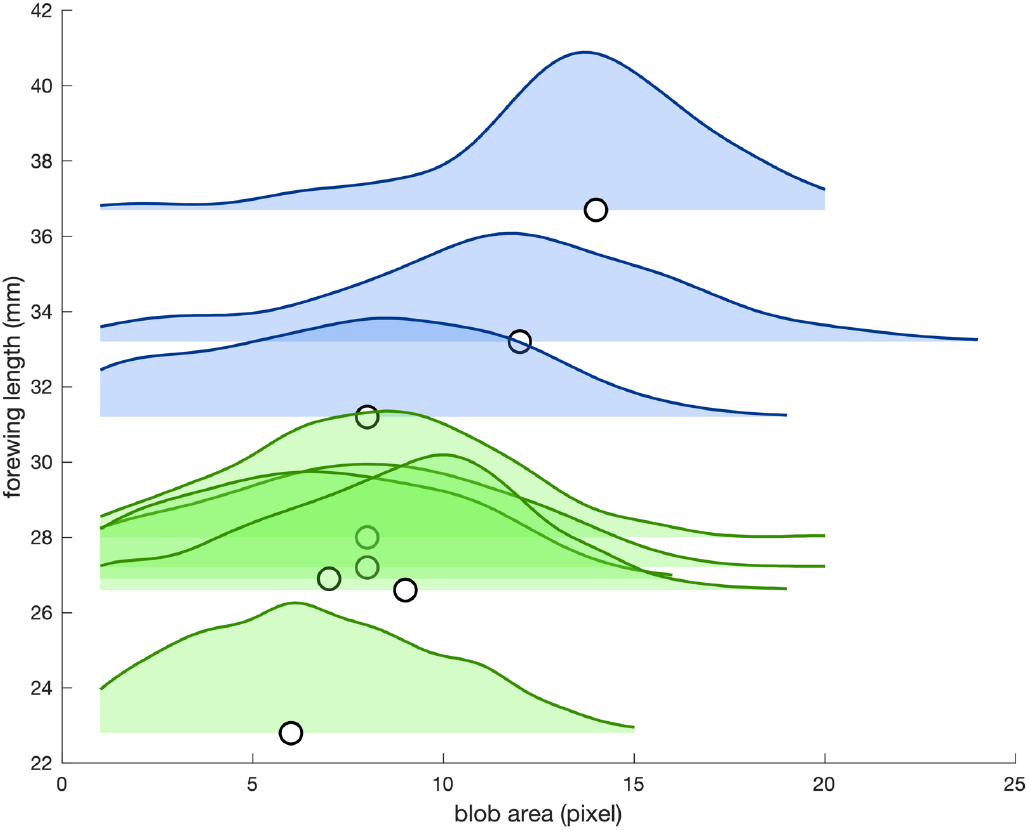
**Forewing length vs. pixel blob area distribution** for 8 tracked and captured *Pieris* butterflies. Blue: *P. brassicae* (*N* = 3); Green: *P. rapae* (*N* = 5). Distributions are displayed as Kernel probability density estimates. Circles indicate the median blob area value for each butterfly.

### Effect of advancing season on flight azimuth

By observing the tracks at different times in the flight period, it appeared visually that the flight azimuths have varied over the season (Fig. 8).

**Figure 8.**
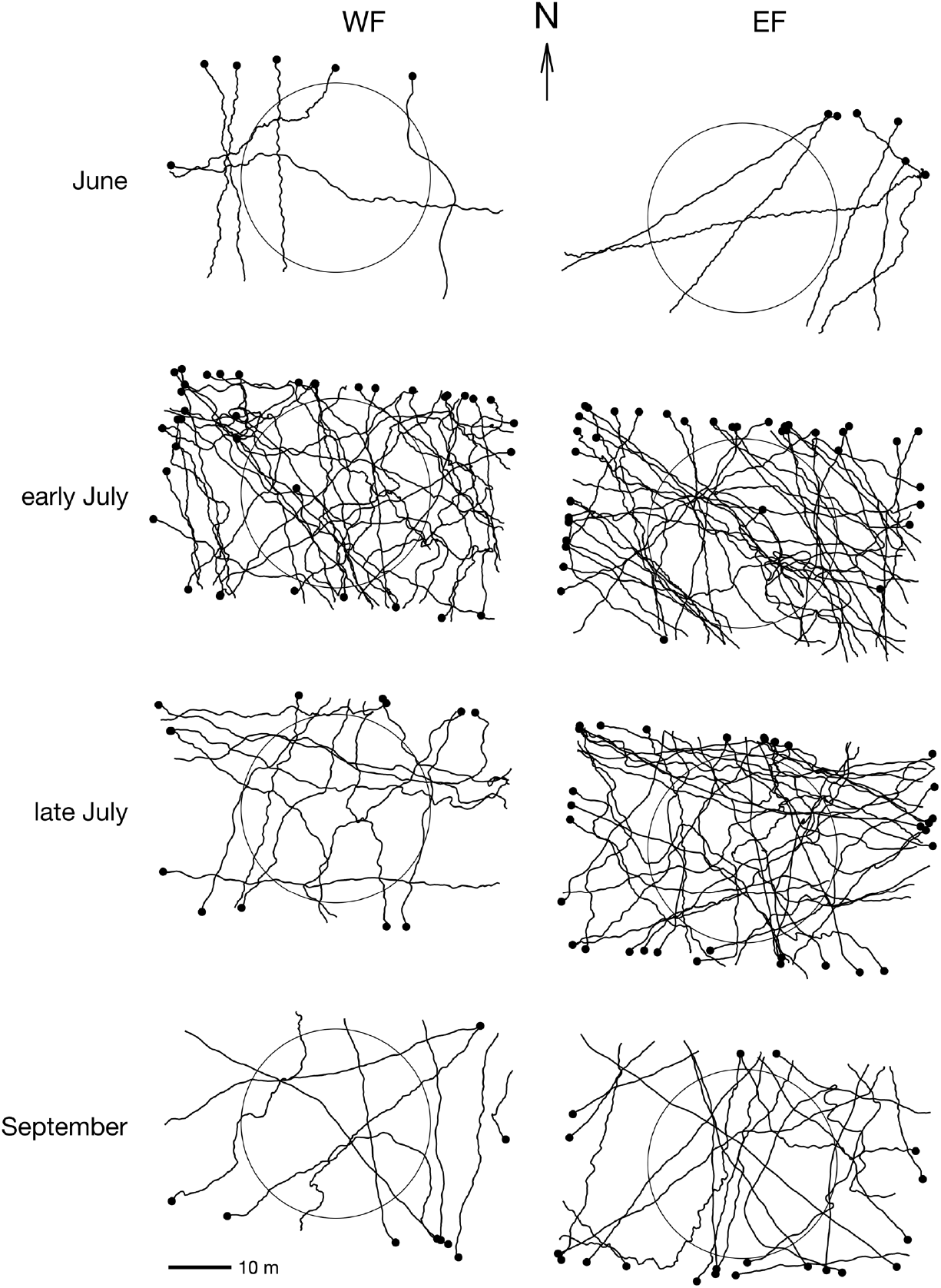
***Pieris* flight tracks broken down by flight period** (vertically) and field (horizontally). Black dots indicate last position of each track (*N* = 166). 30 m diameter circles represent the area considered for azimuth statistical comparisons (*N* = 119).

To further quantify seasonal variations in azimuth distributions, we computed the beeline azimuth of tracks crossing a 30 m diameter disc located at the centre of the camera’s FOV (Fig. 8). Indeed, the rectangular shape of the camera FOV is less likely to record trajectories parallel to the longer side of the FOV (see Fig. S1), and this bias can be corrected by considering only an area enclosed by a circle inside the FOV. When a track did not cross this central disc, it was therefore removed from the sample for circular statistics. When a tortuous track crossed this disc more than once, only the longest segment inside the disc was considered. This restriction of the FOV to a central disc had the effect of reducing our sample from *N* = 166 to *N* = 119 (for this section only).

We tested the effects of period (June to September) and field (WF vs. EF) on beeline azimuth, using a two-factor ANOVA for circular data (Harrison & Kanji 1988, in Berens 2009). As we observed many more trajectories in July, we subdivided the month of July: *early July* (field sessions on July 1^st^ and 8^th^) and *late July* (July 15^th^ and 21^st^). We then tested the uniformity of the azimuth distributions for each period, using Rayleigh tests (Fisher 1995, in Berens 2009).

The ANOVA for circular data detected a significant effect of period on azimuth (*X*^*2*^_(6)_ = 43, *p* = 1.2 × 10^−7^), but no effect of field side (*X*^*2*^_(2)_ = 0.51, *p* = 0.77), and no interaction between the two factors (*X*^*2*^_(3)_ = 5.5, *p* = 0.14). The distribution of azimuths during the 4 periods is shown in Fig. 9. In June, the butterflies flew most often to the north-east (Rayleigh test, *N* = 7, *R* = 0.65, *p* = 0.04; mean azimuth 24°). In early July, they flew most often to the north-west (*N* = 51, *R* = 0.56, *p* = 4 × 10^−8^; mean azimuth 317°). In late July, tracks in all directions were observed, without any dominant azimuth, so that the azimuth distribution was not significantly different from a homogeneous distribution (*N* = 42, *R* = 0.15, *p* = 0.41). Finally, in September, tracks were most often oriented to the south (*N* = 19, *R* = 0.59, *p* = 0.001; mean azimuth 173°).

**Figure 9.**
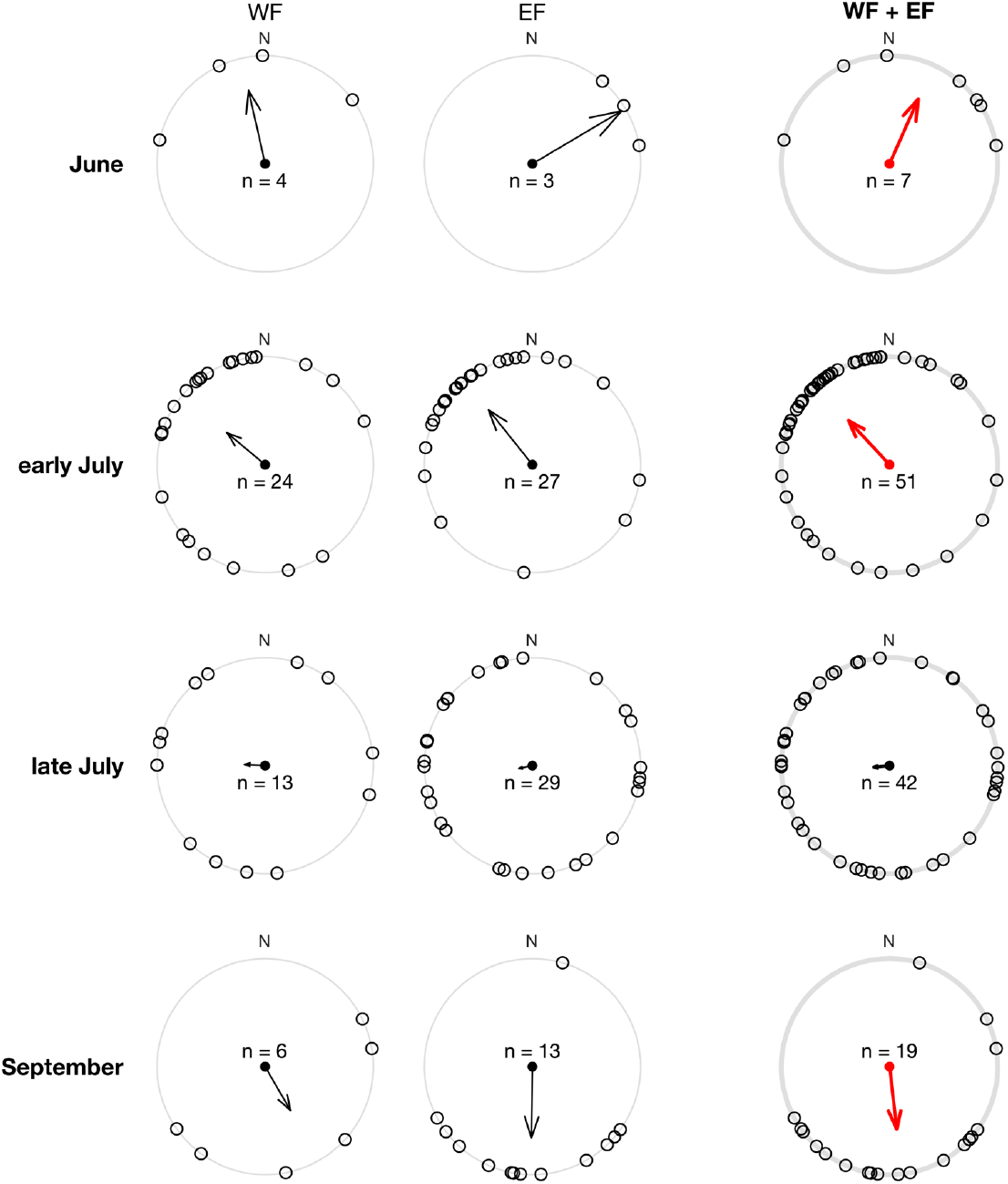
**Circular distributions of *Pieris* beeline azimuth,** broken down by period (vertically), and field (horizontally). Arrows represent mean resultant vectors. The right column shows azimuth distributions for both fields pooled (WF + EF), with red arrows representing significant directional preference according to a Rayleigh test. The Rayleigh test asks how large the mean resultant vector length *R* must be to indicate a non-uniform distribution (Fisher 1995, in Berens 2009).

Although our data was collected in low wind conditions (median wind speed 1.9 m.s^-1^), wind could still influence the butterflies’ trajectory. To verify if the above results were influenced by wind, we first tested whether butterfly beeline azimuth was correlated with wind direction, using a circular-circular correlation test (Jammalamadaka & Sengupta 2001, in Berens 2009).

The test returned no significant correlation (*c* = −0.01, *p* = 0.95; Fig. 10A). We also assessed whether butterfly preferentially flew downwind, crosswind or upwind, by computing the angular difference between beeline azimuth and wind direction, and testing whether this “angle to wind*”* variable departed from a uniform distribution. Results suggested that butterflies did not preferentially fly at a specific angle to wind (Rayleigh test, *N* = 119, *R* = 0.15, *p* = 0.06; Fig. 10B), but the test was close to statistical significance, despite a small resultant vector (i.e. small effect size). Therefore, as a supplementary verification, we focused on tracks recorded during stronger winds (> 2 m.s^-1^, *N* = 64), as these butterflies were expected to be the most affected by a possible wind influence. Both circular-circular correlation (*c* = −0.04, *p* = 0.75) and Rayleigh test on angle to wind (*N* = 64, *R* = 0.08, *p* = 0.67) returned non-significant results (Fig. 10C, D). This comforted the conclusion that wind direction did not significantly bias butterfly flight azimuth in our data.

**Figure 10.**
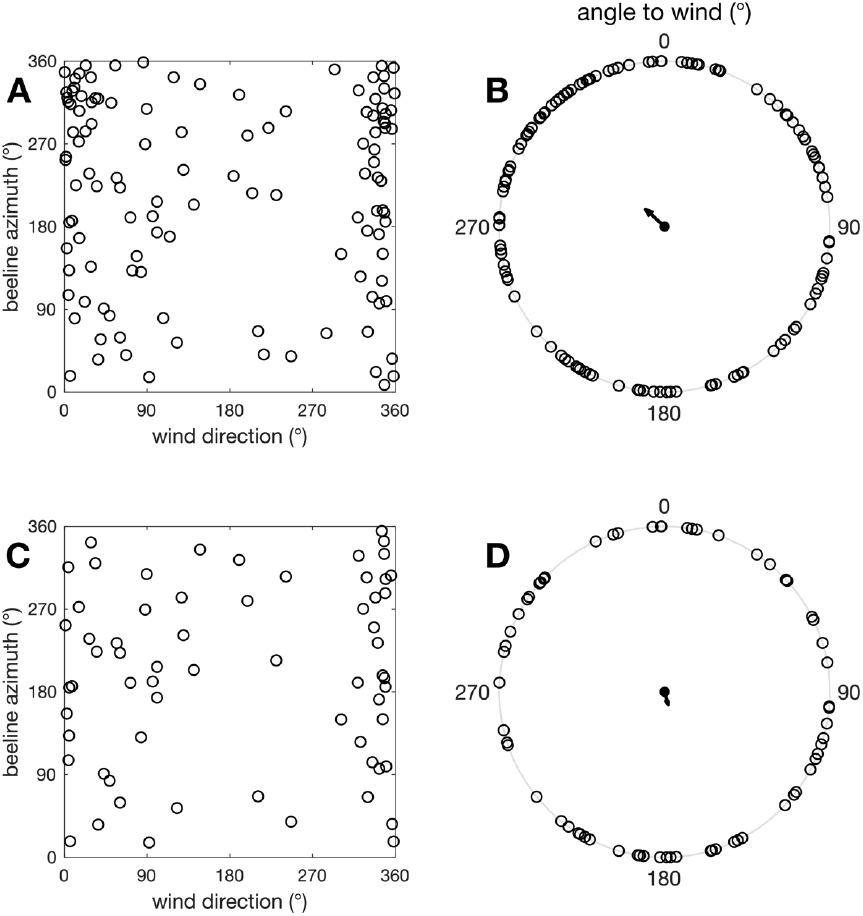
Wind and *Pieris* flight azimuth. **(A)** Beeline azimuth vs. wind direction for *N* = 119 tracks. **(B)** Angle to wind for *N* = 119 tracks. 0° corresponds to upwind flight, 180° to downwind flight. **(C-D)** Same graphics for the *N* = 64 tracks where wind speed exceeded 2 m.s^-1^.

## Discussion

### Butterfly tracking method validation, strengths and limits

We show that UAV-image-based tracking can be used to reconstruct free-flying butterfly paths. With light, affordable and easily-deployed gear in the field, we were able to track numerous wild, untagged *Pieris* butterflies, over an area of 2100 m^2^, with track lengths averaging 40 m. Given an average flight speed of about 3 m.s^-1^, this translates to tracking durations most often near 10-20 s (Fig. 3B-D), depending on each butterfly’s flight speed and straightness. This scale of recorded movement is larger than what can be achieved in most insectaries (e.g. Kleckova et al. 2024: 15 m^2^; Le Roy et al. 2021: 36 m^2^; Kitamura & Imafuku 2015: 182 m^2^; Lihoreau et al 2016b: 880 m^2^), but remains modest compared to other open field methods such as SHR (Ovaskainen et al. 2008: 2.5 km^2^; Lihoreau et al. 2012: 1.5 km^2^; Maggiora et al. 2019: 0.7 km^2^) or human-held GPS tracking (e.g. Fernandez et al. 2016: 100 m average track length).

The spatial scale of the present method is directly constrained by the UAV camera FOV, which was voluntarily limited to 61 × 34 m (from a 45 m UAV height). This was needed to maintain a butterfly blob area around 10 pixels, for reliable blob tracking throughout video frame series (see Methods section). A first possible way to enlarge FOV would be to use an UAV with higher camera resolution (e.g. 6K or 8K sensor, with matching optics quality), which could potentially fly higher and record a wider ground area, while maintaining a centimetric pixel pitch on the ground. Moreover, a higher signal-to-noise ratio in video frames (e.g. from larger sensors and/or less compressed video file formats) might allow to reliably detect smaller blobs throughout frame series, which in turn would allow larger pixel pitch values, and even higher UAV flight height. A FOV exceeding 100 m in side length (i.e. ∼5000 m^2^) is probably already possible with current high-end commercial UAVs, which might be a large enough area to record routine flight movements in some species with limited home ranges (e.g. Fernandez et al 2016).

An alternative way to greatly increase recorded movement length would be to try and follow butterflies with the UAV. This would necessitate (1) that the video feedback to the UAV pilot is of sufficient magnification for a live view of individual butterfly blobs and (2) a different approach to video frame filtering (accounting for quickly moving image background), but these are interesting perspectives for future methodological developments, that might allow recording butterfly movements at a much larger spatial scale, closer to the real scale of dispersal or migration movements.

On the other hand, a strength of the movement data we collected is the 30 Hz temporal resolution, which is orders of magnitude higher than non-image-based field tracking methods applicable to flying insects: 1 Hz for human-held GPS tracking (Fernandez et al. 2016), 0.33 Hz for SHR (Ovaskainen et al. 2008), and lower (usually < 0.01 Hz) for automated radio-tracking (Kays et al. 2011). Here, with one location every ∼10 cm along the flight path, the reconstructed tracks reveal fine-scale movement patterns (see Fig. 2, 4, 8), and offer access to flight speed and tortuosity in the wild, with improved accuracy. Such refined movement data may provide interesting insights on biomechanical and/or orientation processes at work during butterfly flight. More detailed analyses focused on the effect of wind on flight speeds, and the oscillation patterns along *Pieris* flight paths are envisaged, but as they imply many additional analyses (e.g. Srygley & Oliveira 2001), they were beyond the scope of the present UAV methodology presentation. Note that in environments richer in ecological resources (e.g. patches of host plant or nectariferous flowers) and conspecifics, having fine access to flight speed and tortuosity will be useful to study less directed, routine flight behaviours, such as foraging or mate searching (Schtickzelle et al. 2007, Fernandez et al. 2016).

Another limitation of the present method is the projection of the FOV, a pyramidal 3D air volume that butterflies can cross at various height, onto a virtual 2D surface at ground level. For a UAV at height *H* and a butterfly flying at height *h*, this 2D projection at ground level causes an overestimation of flight speed by a factor of *H* / (*H*-*h*). For example, the speed of a butterfly flying at *h* = 15 m under our UAV at *H* = 45 m would be overestimated by a factor of 1.5. In our dataset, we were able to verify by direct observation that at least 27 % of tracked butterflies flew at less than 2 m above ground level. For these tracks, the 2D ground projection implies only a small error on flight speed (overestimation factor ≤ 1.05). For other butterflies, for which we were unable to confirm flight height, we showed that their projected flight speeds are no higher than confirmed “low flyers”, and thus conclude that they probably also crossed the FOV at low height (Fig. 6). This remains indirect evidence, and we cannot exclude that for a small minority of tracks, we might have significantly overestimated flight speed because the butterfly crossed the FOV at a higher height.

Note that, in the hypothetical situation where butterflies move mainly in horizontal planes, but at various heights, the reconstruction of flight speeds will be heterogeneously overestimated, but the angular and temporal variables (e.g. variation of azimuth through time) remain unaffected by flight height.

Beyond the average flight height, all the vertical (z) movements of the butterflies are lost when projected onto a (x, y) horizontal plane. For following studies, it is therefore important to observe the 3D flight behaviour of butterflies beforehand, depending on the species, the type of investigated movement (e.g. foraging, patroling, dispersal or migration), the relief of the terrain and vegetation, and to assess from these necessary preliminary behavioural observations whether a 2D projection might overlook relevant information about the animals’ movements. If the investigated movement is mainly in the horizontal plane and at low height, then the present method can be appropriate. In the case where the 3D flight trajectory or elevation relative to the ground are necessary data for a study, one should either find a zone where the butterfly’s shadow is also visible in the image (Fig. 2H) and derive 3D track data, or opt for other natively 3D optical methods, based on multiple views of the flight volume (Theriault et al. 2014, de Margerie et al. 2015).

When the UAV receives no command from the pilot, its flight control algorithm seeks to maintain its horizontal position, height and azimuth, using sensory-motor regulation loops based on many on-board sensors (GPS, barometric altimeter, magnetic compass, accelerometers, gyroscopes, downward vision system). Commercial UAVs have made spectacular progress on stability in the last decade, and we were able to verify that, at least over the tracking durations used here (< 1 min), the movements of our UAV were indeed very limited (Fig. 5). This allowed us to assume that the drone was static in the sky, at the cost of a drift under 30 cm in the reconstructed position of the butterfly, which we found acceptable for the question posed here (i.e. the measurement of the beeline azimuth over several tens of meters). Still, note that most UAVs use barometric sensors for regulating flight height, and that atmospheric pressure might vary significantly across flight times longer than a few minutes.

For studies requiring lower error caused by the drone’s position, it is possible to perform a continuous measurement of the reference segment on every video frame (by auto-tracking the 2 reference points), or even to perform a more refined calibration of the projected image by continuously tracking multiple points on a grid to fully monitor any complex image transformation or distortion. If the drone’s movements really need to be more closely controlled, more advanced commercial drones are available, using positioning technologies such as Differential GPS (DGPS) or Real Time Kinematic (RTK), which can monitor the UAV position with centimeter accuracy.

Our video tests suggest that other butterfly species are potentially detectable using the present method, whether they appear light against a dark background (resulting in white blobs in filtered video frames) or dark against a light background (black blobs, Fig. 2F). The blink filter proved efficient for erasing background textures, while also tolerating a fair amount of background movement, caused either by the slow drift of the UAV, or vegetation being blown over by the wind. Hence, we hope to see following studies tracking other butterfly species in various open landscapes. In more cluttered landscapes, where the butterflies can fly through or below vegetation, the present optical method will not be appropriate.

Unfortunately, depending on the location and the flight season, it is possible to run into the issue that two (or more) species with similar sizes and colors get filmed simultaneously, and this is the problem we encountered here with two *Pieris* species. This problem of distinguishing species and individuals is often encountered with tagless, image-based or radar-based tracking (Schlagel et al. 2020). We were not able to discriminate *P. brassicae* from *P. rapae* based on pixel blob area (Fig. 7). The wide, overlapping distributions of blob areas for each individual butterfly is not surprising: Flying butterflies can have any posture, from fully stretched to fully closed wings on different video frames, as a result of (1) flapping wing movement and (2) variable roll and pitch angles of the body along the flight path. Moreover, pixel blob area can also be affected by other factors such as (3) contrast with the local background, (4) “blob-clipping” (i.e. when the distance covered between two consecutive frames is less than the body length, which interferes with the filtering method), and (5) flight height. Note also that the relationship between butterfly size and blob area is expected to follow discrete steps (pixels), especially when the pixel pitch is close to animal body size (Fig. 2C).

Provided that some of these sources of blob area variation can be better controlled, we do not rule out that pixel blob area could help discriminate butterfly species in future studies (and/or monitor flight height), but this would need more refined blob classification processes than what we implemented here.

Another option could be to discriminate species based on flight speed or flight behaviour (e.g. tortuosity). A first issue with this approach is that in studies aiming at describing the flight behaviour, it could lead to circular inference. Moreover, Kleckova et al. (2024) reported that different species (*P. rapae* and *P. napi*) can have smaller differences in their flight parameters (measured in an insectary) than spring and summer generations of the same species.

Another route for reducing possible confusion between species could be a greater effort to identify each individual’s species (and sex) in the field, either by remote visual/photographic identification, or by systematically capturing butterflies after they crossed the UAV’s FOV. This would be limited to low-flying butterflies, and also implies greater human presence and movements in the field, which might affect the butterflies’ movement patterns. As well, capturing the butterflies before they cross the FOV, and tracking their movements once released was not retained as a valid option, as released butterflies do not immediately display their normal flight behaviour (Nikoleav 1974, Dudley & Srygley 1994).

### Observed flight speed and straightness distributions

Ground-based multi-camera settings can be used to measure butterfly flight speed in 3D, most often inside insectaries (e.g. Kitamura & Imafuku 2015, Le Roy et al. 2021, Kleckova et al. 2024). Unfortunately, many butterflies species tend to fly slower in insectaries than in the wild (Dudley & Srygley 1994), and captivity can also affect other flight parameters (e.g. glide duration: Le Roy et al. 2021). The present data offers a nice opportunity to measure *Pieris* flight speed in undisturbed, natural conditions, although in 2D only. The median average ground speed we report, at 3.3 m.s^-1^, comes out lower than some earlier measurements on smaller samples of *Pieris* performing directed flight in the field (e.g. 3.6 m.s^-1^ in *P. brassicae*, Nikolaev 1974; 4.4 m.s^-1^ in *P. rapae*, Dunn 2024). More interestingly, across our relatively large sample from multiple field sessions, we observed a wide speed range (from 1.9 to 9.2 m.s^-1^ for track average ground speed, Fig. 3D), not even accounting for flight speed variations within each track. This again calls for a detailled (upcoming) analysis of instantaneaous ground and air speeds, taking wind into account, that might provide novel insights on the flight behaviour of *Pieris* (e.g. wind drift compensation; Gilbert & Singer 1975, Srygley & Oliveira 2001).

Flight speeds over WF (mixed crop) and EF (grass) did not differ, and most tracks had a very high straightness value, compatible with a “directional, undistracted” flight behaviour that is often understood as migratory in butterflies (Chowdhury et al. 2021). Still, we measured that tracks over WF were not quite as straight as over EF (Fig. 3E), suggesting that the richer vegetation in WF might have attracted butterflies to some extent, favouring slightly less directed movement. Also, we note that some butterflies in our sample exhibited clearly tortuous rather than directional flight (see Fig. S2 for 14 tracks with straightness < 0.6, which were observed in equal numbers above WF and EF). Moreover, note that the speed and straightness distributions we report would be different if we had included butterflies marking stops along their flights. Thus, although most *Pieris* butterflies we observed in the field exhibited a directed, possibly migratory behaviour, a minority appeared to be rather engaged in undirected flight movements.

The higher number of butterflies passing over EF (*N* = 100 vs. *N* = 66 for WF) was intriguing. After a close examination of sample sizes for each field session (see table S1), it appears that EF and WF butterfly numbers only differed significantly for 1 out of 11 field sessions, thus we do not conclude that EF consistently attracted more butterflies than WF.

### *P. brassicae* and *P. rapae* migratory behaviours

In butterflies, migration often occurs over several generations, with successive generations following different flight azimuths to achieve a round-trip annual travel (Chowdhury et al. 2021). Within Pieridae, *P. brassicae* and *P. rapae* migratory behaviours have long been documented from the observation of mass migrations (Williams et al. 1942, Vepsalainen 1968). These mass migrations are now rarer due to the use of pesticides (Spieth & Cordes 2012), but see John et al. (2008) and Dunn (2024) for recent reports of group migrations in *P. rapae* (and see Bauer et al. 2024 for radar-based studies on mass migrations in other insect species). Early attempts at quantifying *Pieris* migratory flights used methods such as visually estimating the flight azimuth in the field (e.g. Baker 1968), or difficult mark-recapture experiments (Roer 1959, 1961). More recently, Jones et al (1980) used an egg marking method to study the movements of individual australian *P. rapae* females, reporting directed flight with some northward bias. Gilbert & Raworth (2005) observed in the Pyrenees mountains that a portion of the *P. rapae* population migrated northward in spring and southward in autumn, in line with earlier observations in England (Baker, 1968). For *P. brassicae*, Spieth & Cordes (2012) collected eggs from several Western Europe regions, and later measured the spontaneous flight azimuth of adult female individuals, in a 2 m octogonal flight cage. They showed that the preferred flight azimuth depended on the season and the geographic origin: The first generation usually followed a northward azimuth (modulated by the precise geographic origin), whereas the last generation (2^nd^ or 3^rd^ depending on the region) flew southward. Using a similar flight cage, Larranaga et al. (2013) confirmed a mean northward azimuth in both females and males *P. brassicae* of the first generation.

Here, for a mixed sample of *P. brassicae* and *P. rapae* individuals, we observed flights that were mainly directed northwards in the early season (June, early July), and a southward azimuth in the late season (September). This appears congruent with the existing literature on migration in *P. brassicae* (Spieth & Cordes 2012) and *P. rapae* (Gilbert & Raworth 2005). Hence, despite the limited spatial scale of our movement data, it is probable that the highly directed movements we recorded were segments of migratory flights. Note that the absence of a dominant azimuth for late July might be the result of *P. brassicae* already shifting to southward flight, with *P. rapae* still flying predominantly northward at this period (R. Baker, personal communication).

Tracks recorded from a UAV allow an accurate measurement of flight azimuth, without the need to capture or mark the butterflies (which can affect spontaneous flight behaviour: Nikoleav 1974, Dudley & Srygley 1994), at an intermediate scale between a flight cage (or insectary) and mark-recapture experiments. As a plus, high sampling frequency trajectories contain previously unavailable fine-scale information on flight speeds, tortuosity of the flight path, and patterns of rapid azimuth variation (transverse oscillations). This refined data may be studied in greater detail and has the potential to reveal information on locomotor behaviour and the perceptual mechanisms underlying spatial behaviour. Two major limitations of our approach at this stage are (1) the limited spatial scale and (2) the confusion between *P. rapae* and *P. brassicae*, because of their similar sizes and colors, and their concurrent flight in our geographical area.

## Conclusion

Our results reveal that video tracking of butterflies from a UAV is possible, and capable of providing movement data in fully natural conditions, at an unprecedented spatio-temporal resolution, and at a modest cost. This fine-scale data could prove precious for understanding the spatial behaviour of many butterfly species in open landscapes, and study their movement ecology in various contexts, from routine ressource-searching flights in the local habitat, to dispersal or even migratory flights. We hope that the present methodology exploration can serve as a starting point and motivate other works using UAVs to study spatial behaviour and movement ecology in flying insects.

## Acknowledgements

We thank the owner of the crop fields for authorizing our presence and UAV flights over his property. We thank Robin Baker and Tristan Lafranchis for discussions on the behaviour and biology of *Pieris brassicae* and *Pieris rapae*. We thank Ty Hedrick for discussions on the use of DLTdv. We thank Virginie Durier and Simon Benhamou for discussions on circular statistics. We thank François Cornu and Ignace Cacciaguerra (Pôle drones, CNRS) for their assistance with the UAV pilot certification, and regulations on the use of UAVs in the field. We thank three anonymous reviewers and Nicolas Schtickzelle for their helpful comments on previous versions of the manuscript. Preprint version 5 of this article has been peer-reviewed and recommended by Peer Community In Ecology (https://doi.org/10.24072/pci.ecology.100780; Schtickzelle, 2025).

## Data, scripts, code and supplementary information availability

Data and scripts needed to replicate the statistical results presented here are available at https://dx.doi.org/10.6084/m9.figshare.28043333, as well as supplementary figures and table.

