## Supplementary figures and table for "Tracking butterfly flight in the field from an unmanned aerial vehicle (UAV): a methodological proof of principle"

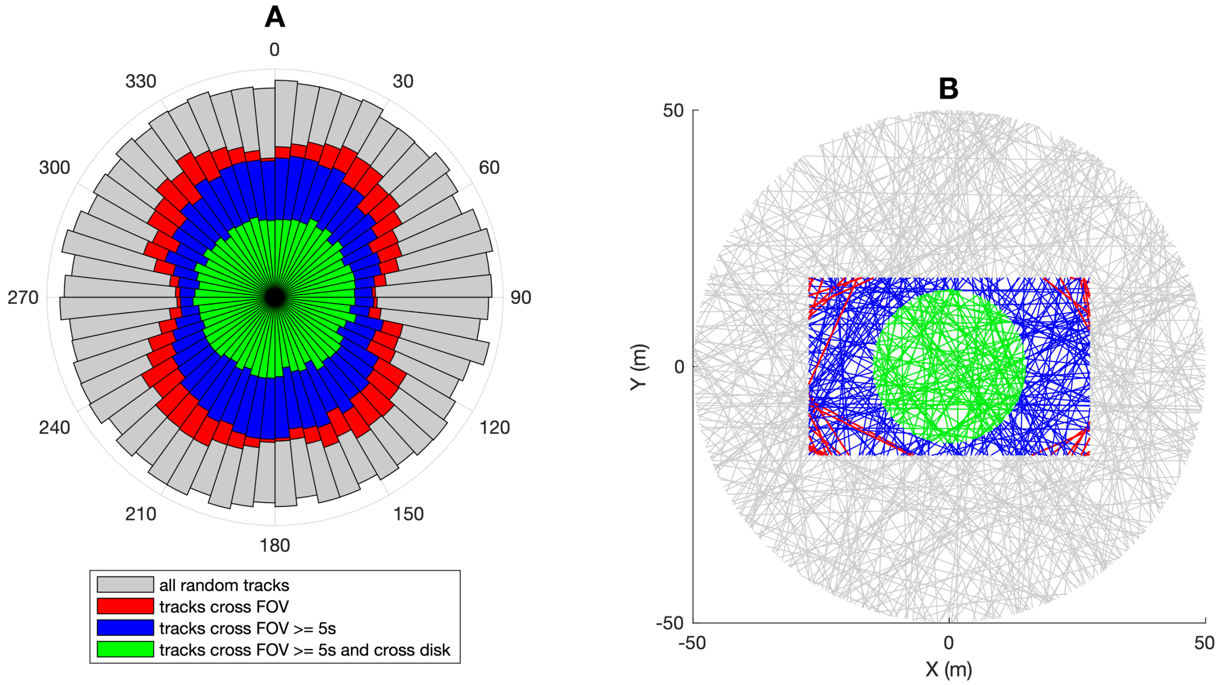


**Figure S1: Azimuth sampling bias caused by a rectangular FOV.**
**A**: The grey polar histogram shows the azimuth distribution of a simulated population of $10^5$ straight tracks in a 2D plane, with random locations and azimuths. Within this population, tracks that cross a rectangular FOV have azimuths that depart from a uniform distribution (red), with oversampling of diagonal directions, and undersampling of directions parallel to the longer side of the FOV. Removing short tracks (< 5 s.) that cross FOV corners reduces oversampling of diagonal directions, but still yields a non-uniform distribution (blue). Only sub-sampling in a central disc (green) within the rectangular FOV reverts to a uniform azimuth distribution. **B**: 2D view of the first 300 tracks in the simulated track population.

*2025-04-02 Version 2*

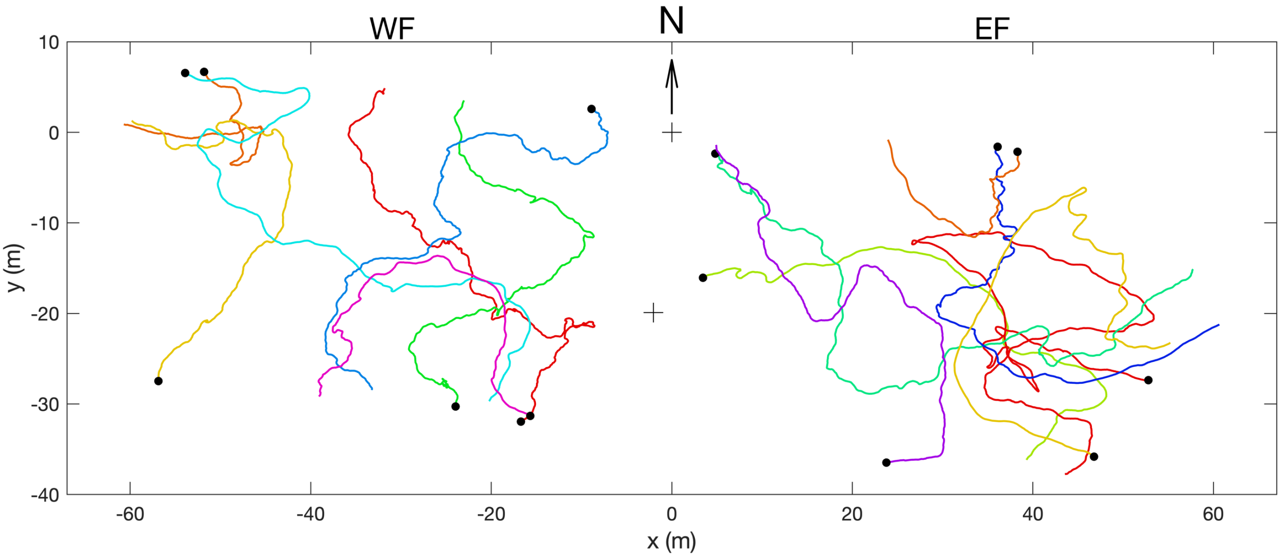


**Figure S2: 2D flight tracks of 14 *Pieris* performing tortuous flight** (straightness < 0.6). Black dots indicate last position of each track. WF: western field; EF: eastern field. These 14 individuals represent the left tail of the distribution in Fig. 3E of the manuscript.

| | | number of tracks | | | | |
|---|---|---|---|---|---|---|
| | field session date | WF | EF | total | proportion of video on WF | p value (binomial test) |
| June | 01/06/2021 | 3 | 0 | 3 | 0,48 | 0,111 |
| | 09/06/2021 | 0 | 0 | 0 | 0,51 | 1,000 |
| | 15/06/2021 | 3 | 6 | 9 | 0,47 | 0,514 |
| early July | 01/07/2021 | 32 | 36 | 68 | 0,48 | 0,504 |
| | 08/07/2021 | 6 | 3 | 9 | 0,50 | 0,508 |
| late July | 15/07/2021 | 8 | 28 | 36 | 0,49 | **0,001** |
| | 21/07/2021 | 5 | 7 | 12 | 0,51 | 0,573 |
| September | 06/09/2021 | 3 | 11 | 14 | 0,48 | 0,059 |
| | 16/09/2021 | 4 | 2 | 6 | 0,51 | 0,688 |
| | 22/09/2021 | 2 | 4 | 6 | 0,47 | 0,691 |
| | 29/09/2021 | 0 | 3 | 3 | 0,47 | 0,252 |
| | all sessions | 66 | 100 | 166 | 0,49 | **0,020** |

**Table S1: Number of reconstructed tracks per field session**, broken down by crop field (WF vs. EF). We compared the observed number of tracks in WF to the expected number based on the proportion of UAV video time spent above WF, using two-tailed binomial tests (Nelson 2015).
